# GATA4 regulates epithelial morphogenesis in the developing mouse stomach to promote establishment of a glandular columnar epithelium

**DOI:** 10.1101/2020.08.18.251587

**Authors:** Ann DeLaForest, Bridget M. Kohlnhofer, Olivia D. Franklin, Roman Stavniichuk, Cayla A. Thompson, Kirthi Pulakanti, Sridhar Rao, Michele A. Battle

**Author notes:** These authors contributed equally to this work (Ann DeLaForest, Bridget M. Kohlnhofer).

## Abstract

The transcription factor GATA4 is broadly expressed in nascent foregut endoderm. As development progresses, GATA4 is lost in the domain giving rise to the stratified squamous epithelium of the esophagus and forestomach (FS), while it is maintained in the domain giving rise to the simple columnar epithelium of the hindstomach (HS). Differential GATA4 expression within these domains coincides with the onset of distinct tissue morphogenetic events, suggesting a role for GATA4 in diversifying foregut endoderm into discrete esophageal/FS and HS tissues. By eliminating GATA4 in the developing HS or maintaining GATA4 in the developing FS, we identified GATA4 as an essential, principal regulator of simple columnar epithelium morphogenesis within the developing HS. GATA4- deficient HS epithelium adopted FS-like fate, and conversely, GATA4- expressing FS epithelium adopted HS-like fate. Underlying structural changes in these epithelia were broad changes in gene expression networks attributable to GATA4 directly activating or repressing expression of HS or FS defining transcripts. Our data implicate GATA4 as having a primary role in suppressing an esophageal/FS transcription factor network during HS development to promote a columnar epithelium. Moreover, GATA4-dependent phenotypes in developmental mutants reflected changes associated with Barrett’s esophagus, suggesting that developmental biology can provide insight into human disease mechanisms.

## Introduction

The mouse stomach consists of two functional domains. The forestomach (FS) is comprised of a keratinized stratified squamous epithelium analogous to the esophagus. The hindstomach (HS) contains a glandular simple columnar epithelium. Although development of these distinct epithelial architectures, each containing specialized cell types, is critical for digestive function, a knowledge gap remains regarding the molecular pathways regulating regionalization. The establishment and maintenance of these domains is crucial for gastrointestinal (GI) tract homeostasis. For example, in Barrett’s esophagus (BE), the stratified squamous epithelium of the distal esophagus converts to a simple columnar epithelium with intestinal and gastric properties (Que et al. 2019). It is clear that defining the molecular mechanisms required for regionalization and maintenance of discrete domains within the GI tract is an essential step towards understanding GI diseases, including BE.

During embryonic development, the GI epithelium emerges from endoderm. Morphogenetic events driven by evolutionarily conserved signaling pathways direct anterior-posterior regionalization of the gut tube into SOX2+ foregut and CDX2+ midgut and hindgut (Thompson et al. 2018). As development progresses, the foregut is further segregated into primitive organ domains, and diversification of the epithelial linings of the esophagus, glandular stomach, and intestines become morphologically apparent. Although not fully understood, specification and differentiation of the epithelium of the esophagus, glandular stomach, and intestines are driven by gene regulatory networks involving lineage restricted transcription factors commonly found to be silenced or amplified in GI diseases, including CDX2, SOX2, p63 and GATA factors 4 and 6 (Thompson et al. 2018; Kim and Shivdasani 2016; Francis et al. 2019; Nekulova et al. 2011; Zheng and Blobel 2010; Sun and Yan 2020; Ayanbule et al. 2011).

The zinc finger containing transcription factor GATA4 is critical for development of many GI organs, including the glandular stomach and small intestine, and is aberrantly expressed in BE and esophageal, gastric, and colorectal cancers (Dulak et al., 2012; Haveri et al., 2008; Jacobsen et al., 2002; Kohlnhofer et al., 2016; Miller et al., 2003; Rodríguez-Seguel et al., 2020; Stavniichuk et al., 2020; Thompson et al., 2017; Walker et al., 2014; Walker et al., 2014b). Between E8.5-E10.5 of mouse development, GATA4 is expressed throughout the caudal foregut endoderm (Jacobsen et al. 2002; Rojas et al. 2010; Watt et al. 2007). By E11.5-12.5, however, GATA4 is lost in the foregut endoderm giving rise to the stratified squamous epithelium of the esophagus and FS, while it is maintained in the foregut endoderm giving rise to the simple columnar epithelium of the HS (Jacobsen et al. 2002). Differential GATA4 expression within these domains further coincides with the onset of divergent tissue morphogenesis suggesting a role for GATA4 in this developmental process. Further supporting a role for GATA4 as a tissue regionalizer is our previous work demonstrating that GATA4 is essential to define the intestinal jejunal/ileal junction (Battle et al., 2008; Thompson et al., 2017). Moreover, histological characterization of chimeric mouse stomachs generated with wild-type and *Gata4^-/-^* ES cells shows that GATA4-null regions display features of stratified epithelium and lose expression of parietal cell, zymogenic chief cell, and neck cell marker genes (Jacobsen et al. 2002). Conditional elimination of GATA4 primarily in the distal region of the mouse stomach during late stages of development using *PDX1-Cre* results in an antrum expressing FS and pancreatic genes (Rodríguez-Seguel et al. 2020). These two mouse models provide insight into the requirement of GATA4 during late stages of glandular stomach development. The role of GATA4 during the earliest stages of foregut development, when domains with different epithelial structures are first delineated, however, has yet to be comprehensively determined.

This study explored the idea that GATA4, acting as a regionalizing factor, is necessary and sufficient to direct development of simple columnar epithelium over stratified squamous epithelium during early gastric development. We used a conditional knockout approach to eliminate GATA4 early in the process of HS development and a conditional knock-in approach to maintain GATA4 expression in the developing FS. We found that GATA4 drives a molecular program in the developing HS by activating expression of a network of genes essential for simple columnar glandular epithelial cell fate while repressing a network of genes essential for stratified epithelial cell identity, including many esophageal-enriched transcription factors. Parallel studies of GATA6 mutants during this developmental window revealed that the phenotype observed in GATA4 mutants was specifically attributable to GATA4 function. Finally, we observed changes in the global transcriptome of GATA4 mutants that overlapped with the metaplastic gene signature of BE suggesting that identifying direct targets of GATA4 in the developing epithelium provides insight into BE.

## Material and Methods

### Mice

The following mouse lines were used: *Gata4^loxP^(Gata4^tm1.1Sad^)*, *Gata4^-^ (Gata4^tmo1Eno^), Shh^tm1^(EGFP/cre)Cjt/J, Gata4^flbio/flbio^*(*Gata4^tm3.1Wtp^)*, *Rosa26^BirA^(Gt(ROSA)26Sor^Tm1(birA)Mejr^), Rosa26 ^lnlG4^ (Gt(ROSA)26Sor ^tm1(Gata4)Bat^, Gata6^loxP^(Gata6^tm2.1Sad^), Gata6^−^(Gata6^tm2.2Sad^)* (Driegen et al. 2005; Harfe et al. 2004; He and Pu 2010; Molkentin et al. 1997; Sodhi et al. 2006; Thompson et al. 2017; Watt et al. 2004). CD1-mice (Charles River Laboratories) were used for control studies and mated with *ShhCre* mice. Timed breeding was used to derive embryos, and a vaginal plug at noon was considered E0.5. Standard PCR protocols were used for genotyping with tail tip, ear punch, or yolk sac DNA. Genotyping primers (Integrated DNA Technologies) are listed in Supplementary Table 1. The Medical College of Wisconsin’s Institutional Animal Care and Use Committee approved all animal procedures.

### Histochemistry and immunohistochemistry

Tissue harvested from the stomachs of E12.5-E18.5 embryos was fixed in 4% paraformaldehyde overnight, dehydrated, paraffin embedded, and sectioned at 5 µm. H&E was performed per standard procedures. Citric acid antigen retrieval was performed for immunohistochemistry with antibodies (Supplementary Table 2). Staining was visualized using R.T.U Vectastain Elite ABC reagent (Vector Labs) and Metal Enhanced DAB substrate kit (ThermoFisher Scientific). Images of histochemical and immunohistochemical stained tissues were obtained either from slides scanned using a NanoZoomer slide scanner and NDP.view 2 software (Hamatsu) or captured using a Nikon Eclipse 80i microscope (Nikon Instruments Inc.) with a Nikon digital smart DS-QiMc camera. Images were assembled into figures using Adobe Photoshop and Illustrator. Images from control and experimental samples were processed identically.

### Quantitative reverse transcriptase polymerase chain reaction

RNA was isolated from epithelial cell preparations from E14.5 and E18.5 embryonic control and mutant FS and HS as previously described (Kohlnhofer et al. 2016). Total RNA was DNase-treated as in previous studies (Bondow et al. 2012; Duncan et al. 1997) or using EZ-DNase (Invitrogen). cDNA was synthesized using MMLV reverse transcriptase (Invitrogen) or using VILO master mix (Invitrogen). TaqMan assays (Supplementary Table 3) were used for qRT-PCR with TaqMan Gene Expression Master Mix (Applied Biosystems). For each gene assayed, three technical replicates were executed on at least three biological samples. *Gapdh* was used for normalization. For each target, expression units were calculated using the formula [2^(-δCq)^]x1000 (Walker et al 2014b). Error bars show standard error of the mean (SEM). Statistical tests employed were either a Student’s T-test or one-way ANOVA, as appropriate.

### RNA-Seq

Total RNA (20 ng) isolated from epithelial cell preparations (Kohlnhofer et al. 2016) was used per biological replicate for RNA-Seq. ERCC RNA Spike-In Controls (ThermoFisher Scientific) were added to each sample prior to Poly-A mRNA selection (New England Biolabs). RNA-Seq libraries were made using the NEBnext Ultra II Directional RNA library prep kit for Illumina (New England Biolabs). The following numbers of embryos were used for RNA-Seq: 3 control FS, 3 control HS, 5 *G4 cKO* HS, and 3 *G4 cKI* FS. All libraries were run as paired-end (38 × 2, total of 76 cycles) on an Illumina NextSeq 500. The raw RNA sequence reads were mapped to mouse reference genome (build mm9) using STARR in Base Pair Tech (basepairtech.com) with default parameters and normalized using the ERCC spike-ins. Quality control matrices were confirmed. Differential expression analysis was completed using the DESeq package. To generate HS-enriched and FS-enriched gene sets, we performed DESeq between FS and HS control data sets and identified enriched genes by p-value ≤ 0.05 and fold change of ≥ 2 criteria. Genes up- or down-regulated in *G4 cKO* HS mutants and *G4 cKI* FS mutants were defined as having a p-value ≤ 0.05 and fold change ≥ 2. Heatmaps were generated using RNA-Seq FPKM values imported into Heatmapper (Babicki et al. 2016).

### ChIP-Seq

Epithelial cells were isolated from adult HS (Thompson et al. 2017) of *Gata4^flbio/flbio^*::*Rosa26^BirA/BirA^* experimental and *Gata4^wt/wt^*::*Rosa26^BirA/BirA^* negative control animals (Driegen et al. 2005; He and Pu 2010). Cells were fixed in 1% formaldehyde for 10 minutes with rocking followed by 5 minutes quenching in 125 mmol/L glycine. Cell pellets were frozen before sonication using a Bioruptor (Diagenode), and GATA4-BioChIP was performed as previously described (Thompson et al. 2017). ChIP-Seq libraries were made using NEBNext Ultra II DNA Library Prep Kit (New England Biolabs). All libraries were run on an Illumina NextSeq 500. The sequencing reads were aligned to the mouse genome (mm9 build) using Bowtie2 using default parameters through Base Pair Tech. Peak calling was done using in-house script employing MACS2 algorithm for all experimental and control samples with q-value cutoff of 0.05 and model fold of 5,50. A total of 50,868 GATA4 peaks were called and were annotated with nearest refseq genes using annotatePeaks.pl script in HOMER v4.11.1 with default settings. The distribution of number of peaks across center of TSS was plotted. Gene targets for peaks in promoters were assigned as the nearest gene (based on TSS position up to +/- 2kb). The genes were imported into DAVID pathway analysis. GO TERM: BP_FAT was reported.

### ChIP-Seq and RNA-Seq Combined Analyses

A list of genes with GATA4 binding peaks located in proximal regulatory regions (+/- 2kb of TSS) was constructed from those genes identified as having HSE or FSE expression as well as differential expression in G*4 cKO* or *G4 cKI* compared with controls. Transcription factor (TF) motifs in this set were identified within +/- 100bp of the center of the GATA4 peak using *findMotifsGenome.pl* script in HOMER v4.11.1 with default settings. HOMER de novo motif discovery strategy searches oligos of 8,10, and 12 bp present in the GATA4 peaks and identifies enriched motifs independent of known TF binding motifs. Enriched oligos are compared with a known TF motif library (JASPAR) to identify predicted matches. The TFs with the closest match to the de novo motif are displayed.

## Results

### GATA4 is essential for simple columnar epithelial development in the mouse hindstomach

GATA4 protein demarcates the stratified squamous FS and simple columnar HS boundary with HS epithelial cells expressing GATA4 (Fig. 1A). We established a *Gata4* conditional knockout (*G4 cKO*) model to determine the extent to which GATA4 is necessary to drive simple columnar epithelial morphogenesis. We compared stomachs across a spectrum of developmental stages among *Gata4^loxP/loxP^*, *Gata4^loxP/+^*, or *Gata4^loxP/-^* controls and *G4 cKO Gata4^loxP/-^::ShhCre* mutant embryos. At E18.5, *Gata4* mRNA was reduced in HS epithelial cells of *G4 cKO* embryos compared with controls (Fig. 1B). Expression of the closely related *Gata6* gene was unchanged (Fig. 1B). H&E staining revealed that the control HS epithelium was appropriately organized into primordial glandular buds lined by simple columnar epithelial cells (Fig. 1C, left). In contrast, the HS epithelial architecture in *G4 cKOs* was severely disrupted (Fig. 1C, middle). Epithelial cells within GATA4-deficient HS lacked columnar morphology, and primordial glandular buds were not present. The mutant epithelium was instead organized similar to a stratified squamous epithelium (Fig. 1C, middle). Reflective of the reduced but not completely extinguished *Gata4* expression in mutant HS (Fig. 1B), we observed that most mutant stomachs contained regions that maintained GATA4 expression, most likely due to Cre inefficiency. Areas with GATA4+ cell patches appeared to have a more columnar, glandular-like morphology (Fig. 1C, right). As expected, FS epithelium of control and *G4 cKO* embryos was phenotypically normal containing keratinized stratified squamous epithelium (Fig. 1D).

**Figure 1:**
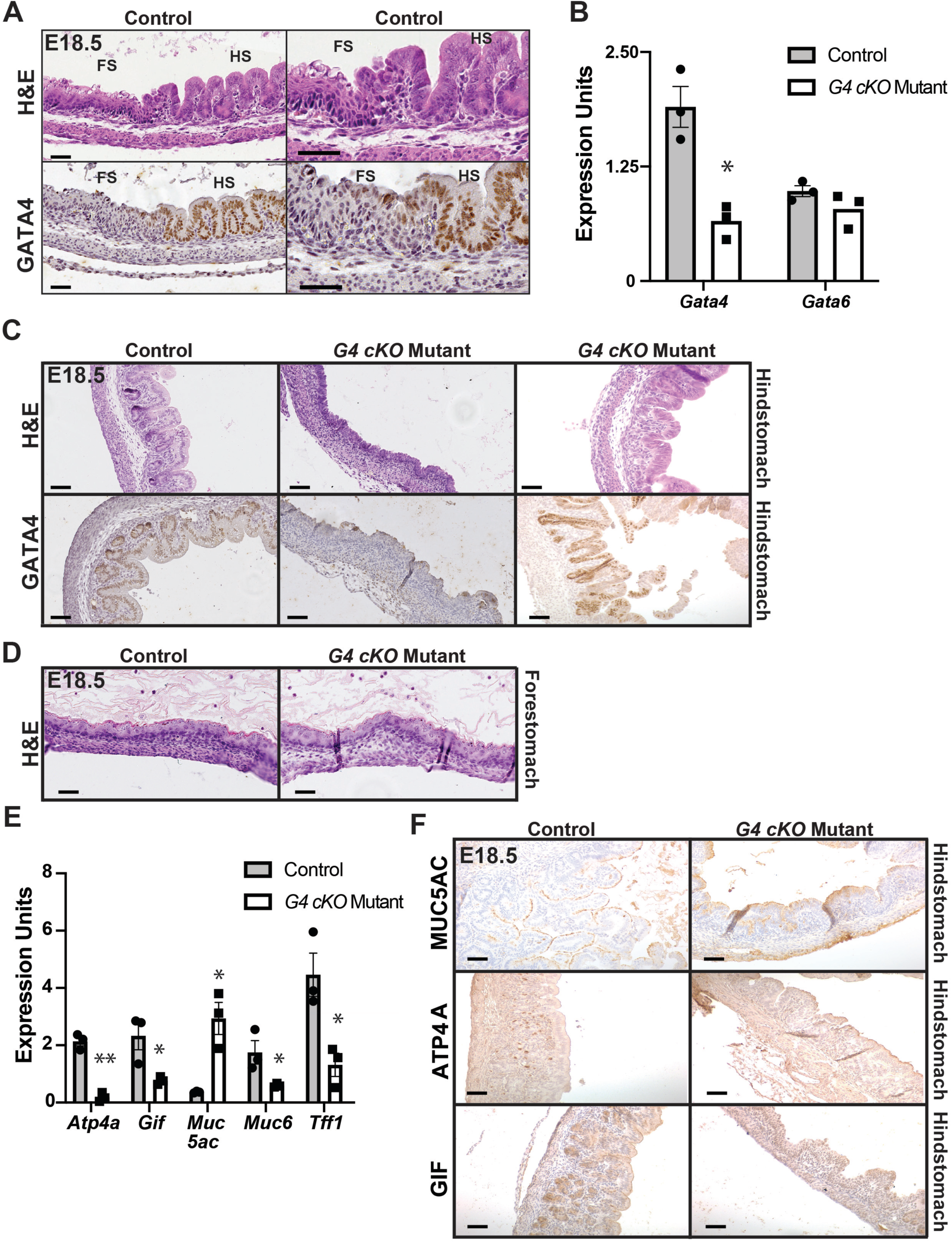
GATA4 is required for morphogenesis of the HS epithelium. (A) H&E and GATA4 antibody staining of E18.5 mouse stomach showed differential GATA4 expression between FS and HS. (B) *Gata4* mRNA was depleted in HS of *G4 cKOs* compared with controls; *Gata6* mRNA remained unchanged (n= 3C,3M). (C) H&E and GATA4 antibody staining of E18.5 mouse control and *G4 cKO* HS (n= 7C,7M) identified GATA4 expressing simple columnar epithelium with emerging gastric units and glands in controls (left). HS of *G4 cKOs* generally lacked GATA4 protein and resembled stratified squamous epithelium (center). Regions in mutants maintaining GATA4 protein were morphologically similar to controls (right). (D) FS from E18.5 control and *G4 cKO* embryos stained with H&E were identical containing keratinized stratified squamous epithelium. (n= 7C,7M) (E) Differentiated HS epithelial cell marker transcripts were changed in *G4 cKO* HS compared with controls (n=3M,3C). All except *Muc5ac* were depleted. *Muc5Ac* increased. (F) MUC5AC protein was unchanged in E18.5 *G4 cKO* HS compared with control (n=4C,4M). ATP4A (n=3C, 3M) and GIF (n=4C,4M) were depleted in in *G4 cKO* HS compared with control. All scale bars, 50μm. All qRT-PCR used isolated HS epithelial cells. Error bars show SEM. P- values determined by a Student’s T-test, \**P*≤ 0.05, \*\**P*≤ 0.001.

We next addressed whether the observed morphological changes in mutant HS reflected changes in the molecular identity of GATA4- deficient cells. To determine the extent to which HS cytodifferentiation was disrupted in E18.5 *G4 cKO* embryos, we measured expression of gastric cell markers in controls and mutants. Transcripts for genes encoding markers of neck (*Muc6, Tff1*), parietal (*Atp4a*), and zymogenic chief cells (*Gif*) were significantly reduced in epithelial cells isolated from *G4 cKO* HS compared with controls; *Muc5ac* (surface mucus cell) transcript was higher in *G4 cKO* HS compared with control (Fig. 1E). MUC5AC (surface mucous/pit cell marker), ATP4A (parietal cell marker) and GIF (zymogenic chief cell marker) proteins were present in control HS, while all but MUC5AC were lacking or diminished in the *G4 cKO* HS (Fig. 1F). These data provide evidence that GATA4 is necessary for HS epithelial morphogenesis and for development of many, but not all, differentiated HS epithelial cell types.

### GATA4 depleted hindstomach acquires expression of genes marking stratified squamous epithelium

The loss of appropriate HS differentiated cell markers paired with the morphological similarity to FS suggested that GATA4-deficient HS epithelium had taken on FS epithelial cell molecular identity. To ascertain if *G4 cKO* HS epithelial cells expressed markers of stratified squamous epithelial cells, we measured transcript and protein levels of key markers of stratified squamous epithelium. We found that transcripts for all FS markers examined—*Krt15, Krt17, Krt23, Cldn10,* and *Trp63*—were ectopically induced in GATA4-deficient HS epithelial cells compared with control cells (Fig. 2A). Induction of FS marker gene expression correlated with ectopic protein expression; KRT5, KRT13, KRT14, and TRP63 were detected in GATA4-deficient HS but not in control HS (Fig. 2B). These data support the idea that GATA4 is necessary for columnar cell development in the HS and that in its absence, abnormal squamous cell development occurs. Notably, *Trp63*, a known master regulator of stratified squamous epithelial cell differentiation was induced in GATA4- deficient HS epithelial cells.

**Figure 2:**
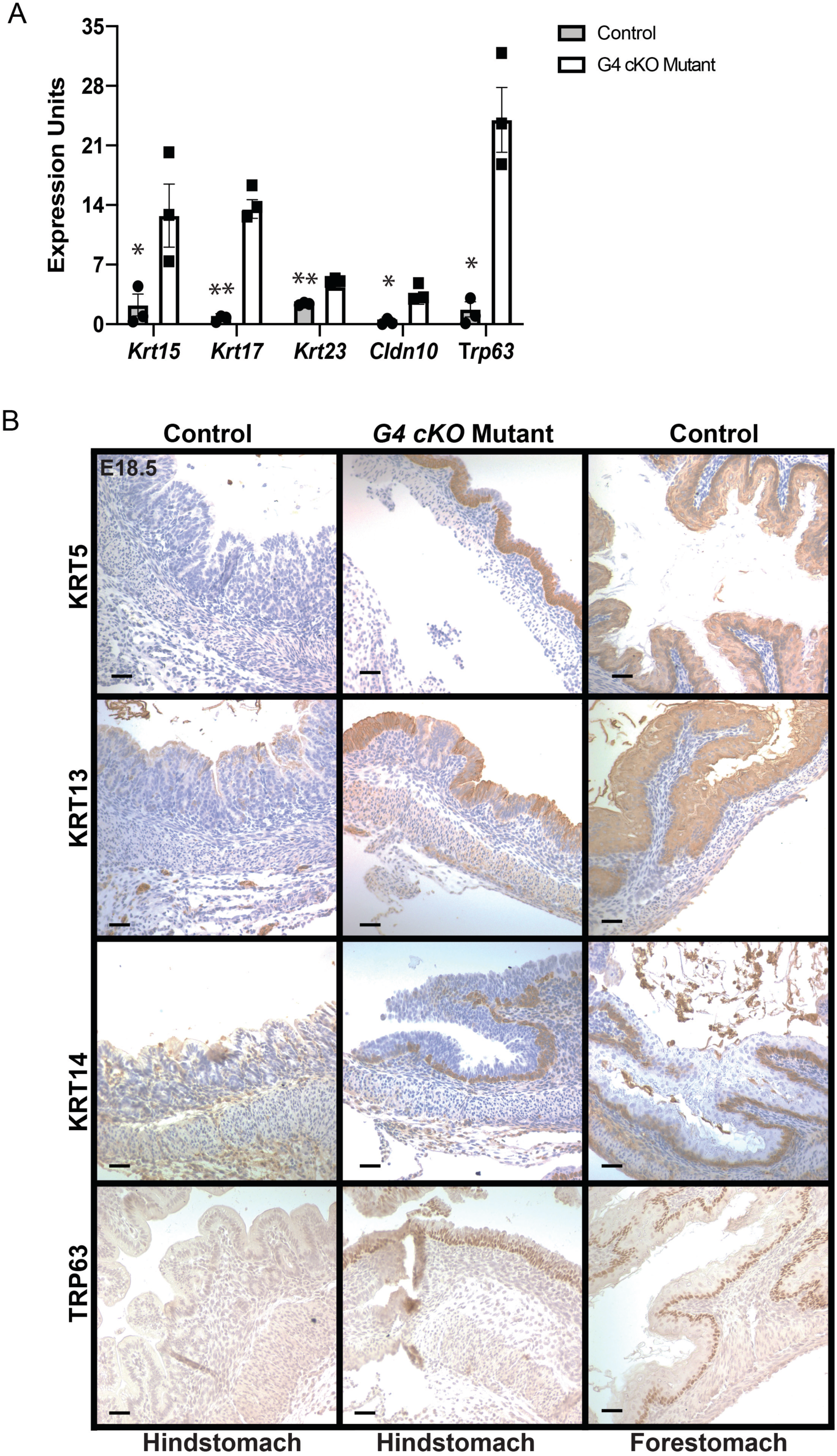
GATA4 depleted HS acquires expression of stratified squamous epithelial cell markers. (A) qRT-PCR with HS epithelial cells from E18.5 controls and *G4 cKOs* showed induced expression of genes associated with stratified squamous epithelium in mutants (n= 3C,3M). Error bars show SEM. P-values determined by a Student’s T-Test, \**P*≤ 0.05, \*\**P*≤ 0.001. (B) FS associated stratified squamous epithelial cell proteins, absent in E18.5 control HS, were induced in *G4 cKO* HS. FS expressed all proteins. (KRT5, n=3C, 5M; KRT13, n=6C, 5M; KRT14, n=4C, n=5M; TRP63 n= 5C, 5M) Scale bars, 50μm.

### Developmental morphological milestones are absent in GATA4 cKO hindstomach by E14.5

Because ShhCre drives gene deletion as early as E8.5-E9.5, we traced back GATA4 depletion in the HS of *G4 cKO* embryos to identify the first time point at which both GATA4 was depleted and phenotypic changes were observable (Fig. 3A-P). At E16.5, GATA4 loss was apparent in mutants, and the GATA4-deficient epithelium was disorganized lacking primordial gastric glands and appearing mainly stratified (Fig. 3A,B,I,J). Similar to what we observed at E18.5, GATA4+ cell patches were present in E16.5 *G4 cKO* HS tissue, and the morphology of these regions more closely resembled the normal emerging columnar, glandular epithelium (Fig. 3A,B,I,J). At E14.5, disrupted GATA4 expression was observed in mutant HS (Fig. 3C,D). HS epithelium of controls displayed an increased apical-basal thickness (compared with E12.5 stomachs) and intraepithelial spaces were present, which have been identified as a precursor to gland and gastric unit development (Fig. 3K,L) (Karam et al. 1997). In contrast, mutant *G4 cKO* HS epithelial thickness was reduced compared with controls, and there were fewer, smaller intraepithelial spaces (Fig. 3K,L). Again, in regions of GATA4 positivity, HS architecture more closely resembled that of control tissue (Fig. 3E,F,M,N). Finally, although GATA4 protein was reduced in HS at E12.5 (Fig. 3G,H), the epithelial HS morphology of controls and mutants was comparable (Fig. 3O,P). Based on these data, we concluded that E14.5 represented the first timepoint that depletion of GATA4 in the developing HS correlated with phenotypic changes. Therefore, we performed subsequent analyses with tissues from E14.5 embryos.

**Figure 3:**
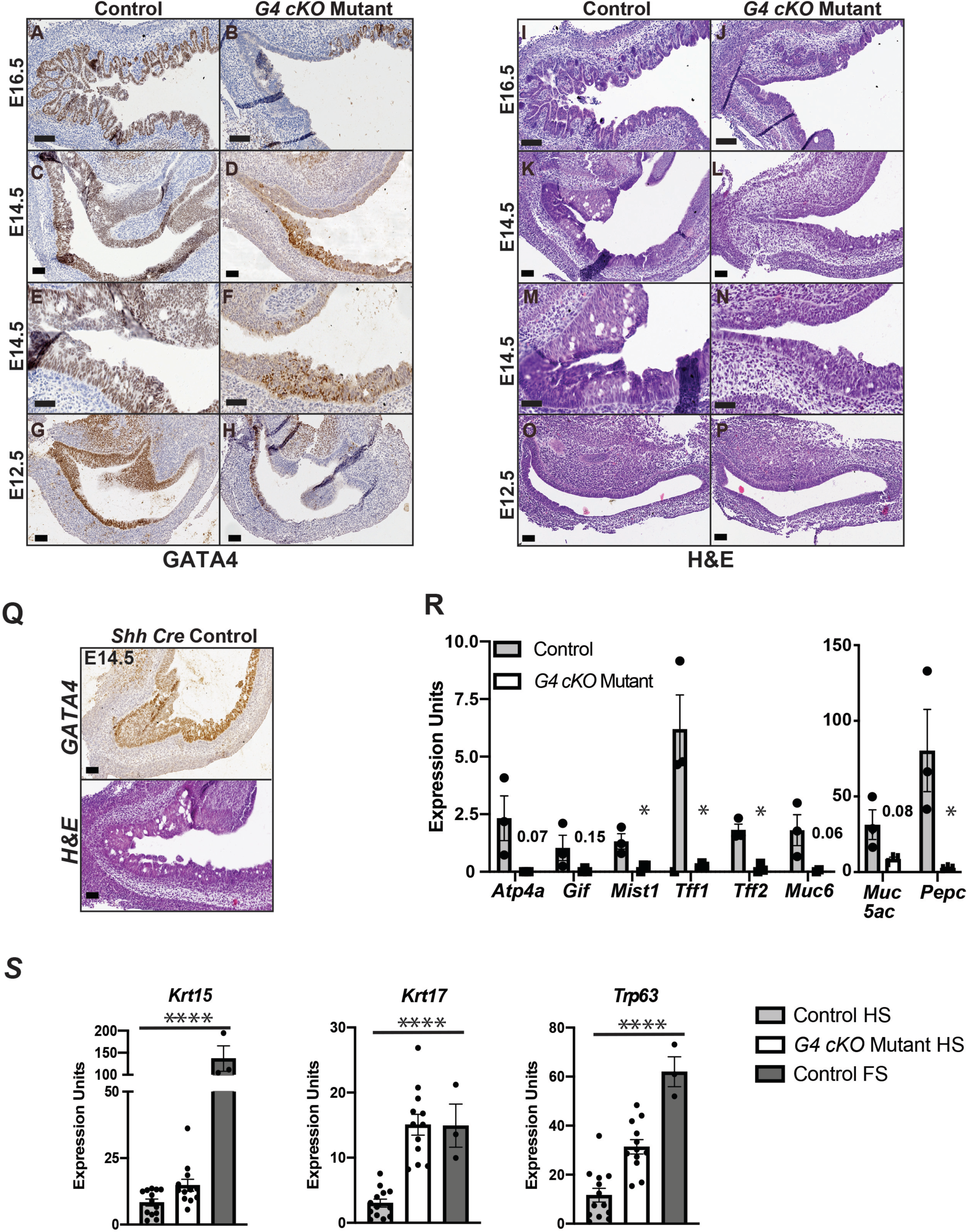
GATA4-deficient HS architecture and gene expression are disrupted at E14.5. (A-H) GATA4 IHC of control and *G4 cKO* HS from E16.5 (n= 6C, 6M), E14.5 (n=7C, 11M) and E12.5 (n= 5C, 6M) embryos showed depleted GATA4 protein in mutants at all stages. Small regions of GATA4+ cells were evident in mutant HS. GATA4 protein loss became more evident in mutants between E12.5 to E14.5. (I-P) H&E staining of control and *G4 cKO* HS from E16.5 (n= 6C, 6M), E14.5 (n=5C, 8M) and E12.5 (n= 4C, 4M) embryos showed disrupted epithelial structure at E16.5 and E14.5. E16.5 control HS epithelium was columnar and lined with primordial buds whereas *G4 cKO* HS epithelium appeared stratified, containing few abnormal developed primordial buds. E14.5 control HS epithelium had a typical morphology containing intraepithelial spaces, which are associated with glandular morphogenesis, whereas mutant HS epithelium was thinner with sparse, smaller intraepithelial spaces. (Q) GATA4 protein expression and epithelial structure were normal in HS from *Gata4^+/+^Shh Cre* E14.5 embryos (n=2) (R) Differentiated HS epithelial cell marker transcripts were decreased in E14.5 *G4 cKO* HS compared with controls (n=3M,3C). (S) Expression of stratified squamous epithelial cell transcripts was increased in E14.5 *G4 cKO* HS compared with controls. All were expressed in control FS (n=13C HS, 12M HS, 3C FS). All scale bars, 50μm. All qRT-PCR used isolated HS or FS epithelial cells. Error bars show SEM. P-values determined by a Student’s T-test, \**P*≤ 0.05, \*\**P*≤ 0.001 or one-way ANOVA as appropriate. \**P*≤ 0.05, **P≤ 0.001, \*\*\*\**P*≤ 0.0001.

To confirm that changes observed were independent of ShhCre, we examined GATA4 protein levels and epithelial morphology of HS from *Gata4^+/+^ShhCre* E14.5 embryos and found these to be indistinguishable from control HS, implying that disruption of one *Shh* allele by Cre insertion did not affect HS development (Fig. 3Q). To examine the extent of cytodifferentiation in E14.5 *G4 cKO* HS, we measured expression of genes marking specific glandular epithelial cell types. We detected decreased levels of parietal cell (*Atp4a, PepC*), zymogenic chief cell (*Gif, Mist1*), neck cell (*Tff1, Tff2, Muc6*), and surface mucous cell (*Muc5Ac*) transcripts in *G4 cKO* HS epithelial cells compared with controls (Fig. 3R). Furthermore, to determine if FS-specific gene expression was induced in the HS epithelium of E14.5 *G4 cKO*s, we performed qRT-PCR for *Krt15*, *Krt17,* and *Trp63* and found these to be ectopically expressed in GATA4- deficient HS epithelium compared with control (Fig. 3S).

### Loss of GATA6 in the hindstomach does not disrupt epithelial morphogenesis

Both GATA4 and GATA6 are expressed in the developing HS and absent in FS (Fig. 1A, 4A). We considered the possibility that phenotypes observed in GATA4 HS mutants reflected an imbalance of total GATA protein dosage rather than loss of GATA4 itself. Therefore, we examined HS from control *Gata6^loxP/+^* and mutant *Gata6^loxP/-^::ShhCre* embryos (*G6 cKO*) at E18.5 and E14.5. At E18.5, steady-state mRNA levels of *Gata4* were similar between control and *G6 cKO* HS epithelial cells, and, as expected, *Gata6* mRNA was depleted in *G6 cKO* HS epithelial cells (Fig. 4B). GATA4 and GATA6 proteins were present in control HS epithelial cells, whereas, only GATA4 protein was retained in the *G6 cKO* mutant HS epithelium (Fig. 4C). H&E staining showed that epithelial architecture was comparable between genotypes. Steady-state mRNA levels of HS cell markers were assessed similarly as was done for GATA4 HS mutants, and no expression changes were detected in *G6 cKO* HS epithelial cells compared with control (Fig. 4D). Finally, we validated that GATA6 protein was efficiently deleted at E14.5 as in GATA4 mutants (Fig. 4E). GATA4 protein was present throughout control and *G6 cKO* HS whereas GATA6 protein was depleted in *G6 cKO* HS. Therefore, we concluded that the phenotypes evident in *G4 cKO* HS were specific to GATA4 deletion rather than an overall reduction of GATA protein.

**Figure 4:**
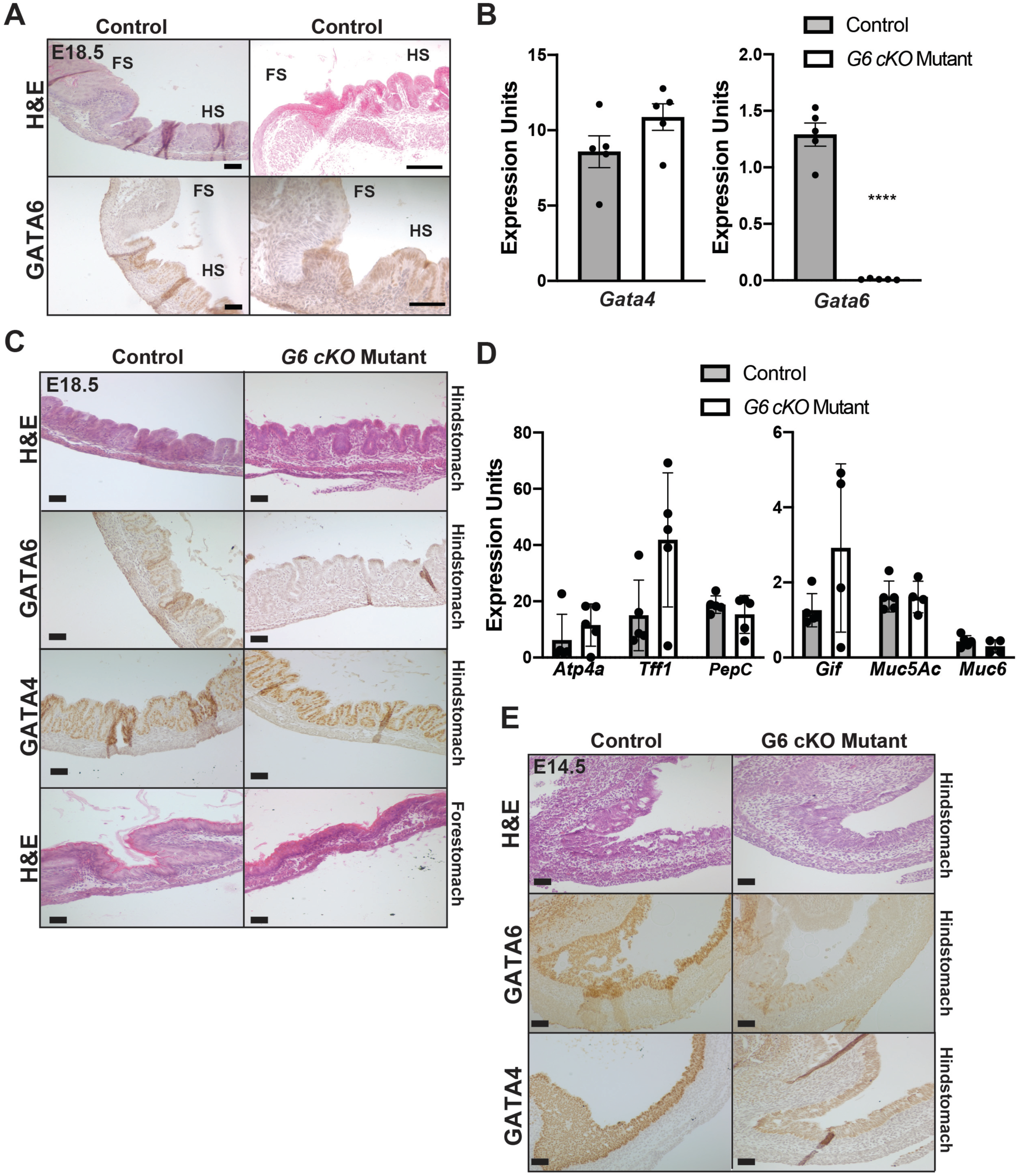
GATA6-depleted HS epithelium develops normally. (A) H&E and GATA4 antibody staining of E18.5 mouse stomach showed differential GATA6 expression between HS and FS. (B) *Gata6* mRNA was depleted in HS of *G4 cKOs* compared with controls; *Gata4* mRNA remained unchanged (n= 5C,5M). (C) H&E GATA6, and GATA4 staining of E18.5 HS control and *G6 cKO* HS showed GATA6 protein depletion in mutants yet epithelial structure remained normal and GATA4 protein was unchanged. FS morphology (H&E staining) was comparable between controls and mutants (n=4C, 4M). (D) Differentiated HS epithelial cell marker transcripts were unchanged in *G6 cKO* HS compared with controls (n=5M,5C). (E) H&E, GATA4, GATA6 staining of control and *G6 cKO* HS at E14.5 (n=6C, 6M) showed comparable morphology between tissues with GATA6 protein depleted and GATA4 protein unchanged. All scale bars, 50μm. All qRT-PCR used isolated HS epithelial cells. Error bars show SEM. P-values determined by a Student’s T-test, \*\*\*\**P* ≤ 0.0001.

### Induction of GATA4 expression in the developing forestomach epithelium alters epithelial architecture toward columnar

Observing that GATA4 expression was necessary to drive columnar cell fate in the developing HS epithelium, we sought to determine whether GATA4 was sufficient to induce columnar fate in the developing FS epithelium. Using *ShhCre* to drive expression of a *Gata4* conditional knock-in allele, *Rosa26 ^lnlG4^* (*Gt(ROSA)26Sor ^tm1(Gata4)Bat^, G4 cKI*) (Thompson et al. 2017) in the developing FS, we analyzed the effects of ectopic GATA4 expression on FS epithelial morphogenesis at E14.5, parallel to studies of *G4 cKO* mutants. We detected *Gata4* mRNA in E14.5 *G4 cKI* FS epithelial cells (Fig. 5A). Compared with GATA4 levels in E14.5 HS, levels in *G4 cKIs* were ∼3-fold lower. Ectopic GATA4 expression in FS, even at a level lower than HS, was sufficient to inhibit expression of *Trp63* transcript (Fig. 5A). This was the inverse of what occurred in *G4 cKO* HS mutants in which *Trp63* expression was induced upon GATA4 depletion (Fig. 2A). The finding that a key marker of stratified squamous epithelial cells was reduced in *G4 cKI* FS suggested a change in cell identity. Therefore, as a preliminary examination of the emergence of HS gene expression in mutant FS epithelium, we measured expression of *Hnf4α,* a factor specifically present in developing HS but absent in developing FS. We detected *Hnf4α* mRNA in *G4 cKI* FS but not in control FS (Fig. 5A). As expected, GATA4 protein was undetectable in E14.5 control FS epithelium whereas FS epithelial cells of *G4 cKI* embryos uniformly expressed GATA4 protein (Fig. 5B). Examination of the epithelial architecture of control and GATA4-expressing FS indicated that control epithelium was developing as a stratified epithelium while the GATA4-expressing FS epithelium appeared columnar-like (Fig. 5B). To further probe the character of the developing GATA4-expressing FS epithelium, we used IHC to compare expression of the stratified squamous cell cytokeratin KRT5 between controls and mutants. Although present in control FS epithelium, KRT5 protein was uniformly depleted in GATA4- expressing FS epithelium (Fig. 5B). These data provide evidence that GATA4 protein is sufficient to drive a columnar epithelial fate in the developing murine stomach.

**Figure 5:**
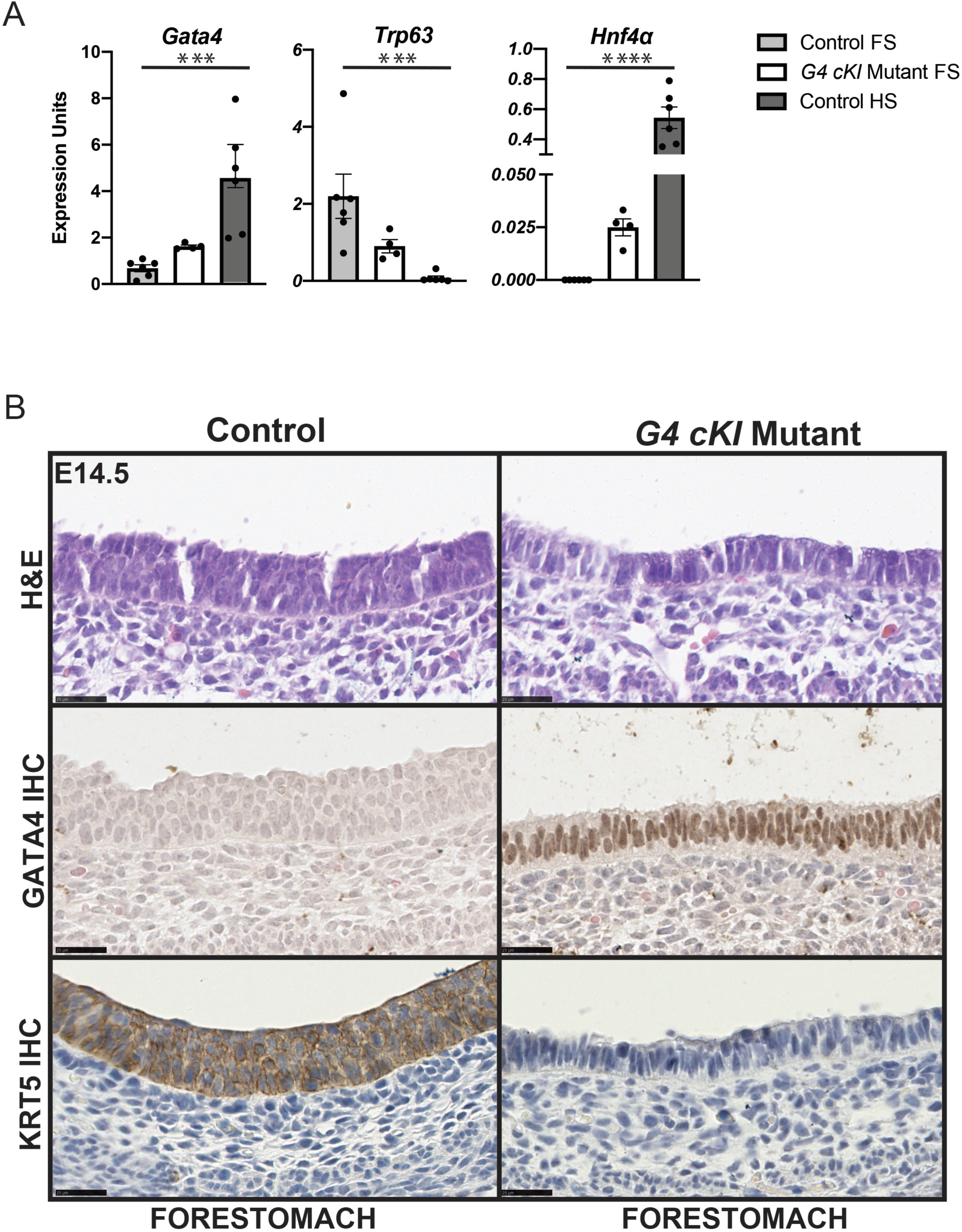
Maintained GATA4 expression in the developing FS epithelium alters epithelial architecture toward columnar. (A) *Gata4, Trp63,* and *Hnf4a* mRNA levels were measured in epithelial cells from control FS, HS, and *G4 cKI* FS at E14.5 (n=6C FS and HS, 4M FS). *Gata4* was up-regulated in *G4 cKI* FS. *Trp63* was highly expressed in control FS and diminished in *G4 cKI* FS. *Hnf4a* was induced in *G4 cKI* FS. Error bars show SEM. P-values were calculated using a one-way ANOVA, \*\*\**P* ≤ 0.005, \*\*\*\**P* ≤ 0.0001. (B) H&E, GATA4, and KRT5 staining of E14.5 control and *G4 cKI* FS (n=3C, n=3M). Control FS contained a multi-layer epithelium whereas *G4 cKI* FS appeared columnar. GATA4 protein was ectopically expressed in E14.5 *G4 cKI* FS epithelial cells. Control FS epithelial cells expressed KRT5 protein that was completely absent in FS of *G4 cKI*s. Scale bars, 25μm.

### The transcriptome of GATA4 gastric mutants reflects global shifts in epithelial cell identities

Our morphological analyses of GATA4-depleted HS epithelium and GATA4-expressing FS epithelium support the idea that GATA4 is necessary and sufficient to promote development of columnar epithelium in the embryonic stomach. Given that GATA4 has been shown to be a powerful transcription factor in regulating jejunal-ileal boundaries and cell fates in the small intestine, we reasoned that global shifts in the overall transcriptomes of these developing gastric tissues must be occurring when GATA4 was modulated (Kohlnhofer et al. 2016; Thompson et al. 2017). Therefore, we performed RNA-Seq using HS and FS epithelial cells from control and mutant E14.5 embryos. We used gene expression profiles of control FS and HS to construct HS enriched (HSE) and FS enriched (HSE) gene sets by selecting genes with ≥ 2-fold differential gene expression (*P*≤ 0.05) between HS and FS (Fig. 6A). We found 1461 genes encoding HSE transcripts and 2619 genes encoding FSE transcripts (Fig. 6A). We compared the sets of genes differentially expressed between control HS and GATA4-depleted HS or control FS and GATA4-expressing FS with HSE and FSE gene sets. Of the genes differentially expressed in *G4 cKO* HS epithelial cells versus control, 46% overlapped with the HSE and FSE gene sets (Fig. 6B). Of the genes differentially expressed in *G4 cKI* FS epithelial cells versus control, 29% overlapped with the HSE and FSE gene sets (Fig. 6C). Reflecting the HS to FS transition in GATA4-deficient HS mutants, 95% of genes overlapping with the HSE set were down-regulated (519/544), and 93% of genes overlapping with the FSE set were up-regulated (1155/1237) (Fig. 6B). Similarly, in GATA4-expressing FS mutants, in which FS transformed to HS, 95% of genes overlapping with the FSE set were down-regulated (737/779), and 80% of genes overlapping with the HSE set were up-regulated (274/363) (Fig. 6C). We further used gene set enrichment analysis (GSEA) to compare control and mutant gene expression profiles. GSEA provides an unbiased computational method to determine whether a defined gene set (HSE or FSE) shows significant enrichment in 1 of 2 biological states (HS epithelium of *G4 cKO* vs control embryos or FS epithelium of *G4 cKI* vs control embryos) (Subramanian et al. 2005). We performed GSEA using HSE (1461) and FSE (2619) gene sets (Fig. 6A) and RNA-Seq data profiling gene expression of HS epithelial cells from *G4 cKO* and control embryos and of FS epithelial cells from *G4 cKI* and control embryos. The global gene expression profile of *G4 cKO* HS epithelium, unlike the control profile, aligned with the FSE gene set (Fig. 6D, top panel). Similarly, the global gene expression profile of *G4 cKI* FS epithelium, unlike the control profile, aligned with the HSE gene set (Fig. 6D, bottom panel). These data further support the idea that GATA4 is necessary and sufficient to promote development of columnar epithelium in the embryonic stomach. Loss of GATA4 in the HS epithelium causes a columnar-to-stratified switch and gain of GATA4 in the FS epithelium causes a stratified-to-columnar switch. Moreover, changes are not simply morphological in nature, but represent global shifts in foundational gene expression patterns that underlie columnar versus stratified epithelial morphogenesis in the developing stomach.

**Figure 6:**
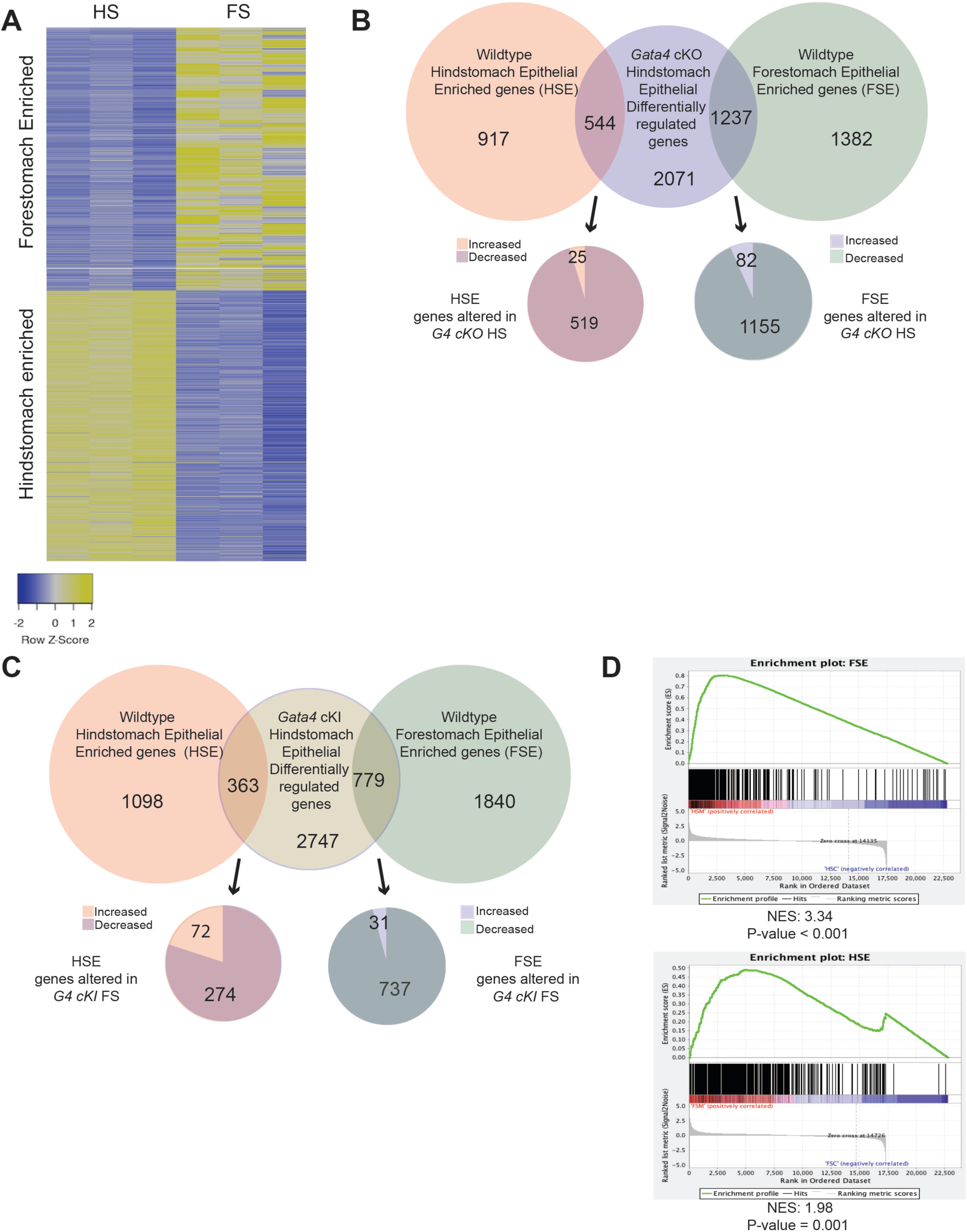
GATA4 mutant HS and FS epithelial cells undergo broad shifts in gene expression correlating with morphology. (A) HS-enriched (HSE) and FS-enriched (FSE) epithelial cell gene expression patterns were determined by comparing gene expression (FPKM) between E14.5 control HS and FS (RNA- Seq, n=3C HS,3C FS). Criteria were ≥ 2-fold expression change and *P*≤ 0.05. Heatmap displays FPKM for HSE and FSE gene sets. (B) Genes differentially expressed between E14.5 *G4 cKO* HS and control HS epithelial cells were identified using criteria ≥ 2-fold expression change and *P*≤ 0.05 (n=3C, n=5M). These genes were overlaid with HSE and FSE sets defined in (A). Overlapping genes were discriminated by direction of gene expression change. Most differentially expressed HSE genes were down-regulated in mutant tissue and most differentially expressed FSE genes were up-regulated in mutant tissue corresponding with observed morphological changes. (C) Differentially expressed genes between in E14.5 *G4 cKI* FS and control FS epithelial cells mapping to HSE or FSE gene sets were determined as in (B, n=3C, n=3M). Most differentially expressed FSE genes were down-regulated in mutant tissue and most differentially expressed HSE genes were up-regulated in mutant tissue corresponding with observed morphological changes. (D) Gene Set Enrichment Analysis performed using HSE and FSE gene sets and differentially regulated genes in *G4 cKO* and *G4 cKI* E14.5 epithelial cells indicated the *G4 cKO* HS epithelial cell gene expression aligned more closely with the FS while *G4 cKI* FS epithelial cell gene expression aligned more closely with the HS. Differentially expressed gene lists can be found in supplemental table 4.

### GATA4 likely directly activates and represses gene expression to control epithelial morphogenesis in the developing hindstomach

Our next goal was to identify genes among those altered in GATA4- deficient HS and GATA4-expressing FS most likely to be directly regulated by GATA4 to establish a simple columnar epithelial cell fate and repress a stratified squamous epithelial cell fate. We took advantage of GATA4-bio-ChIP-Seq data that we generated using control adult mouse HS epithelial cells to identify those genes differentially expressed between controls and mutants that also contained experimentally validated GATA4 binding sites. We identified 50,869 total GATA4 binding peaks in HS epithelium with 41,332 peaks within +/-50 kb of the nearest TSS and 18,658 peaks within +/- 2 kb of the nearest TSS (Fig. 7A). We used DAVID functional annotation to identify enriched biological themes within the identified GATA4 binding data and found themes including epithelium development and morphogenesis, columnar epithelial cell differentiation, and glandular epithelial cell development (Fig. 7B, upper panel), agreeing with our phenotypic data showing that GATA4 promotes columnar cell development. Analysis of the DNA sequence of GATA4 ChIP-Seq peaks within +/- 2kb of the nearest TSS using HOMER showed that the top motif identified matched the GATA4 consensus motif (Fig. 2B, bottom panel).

**Figure 7:**
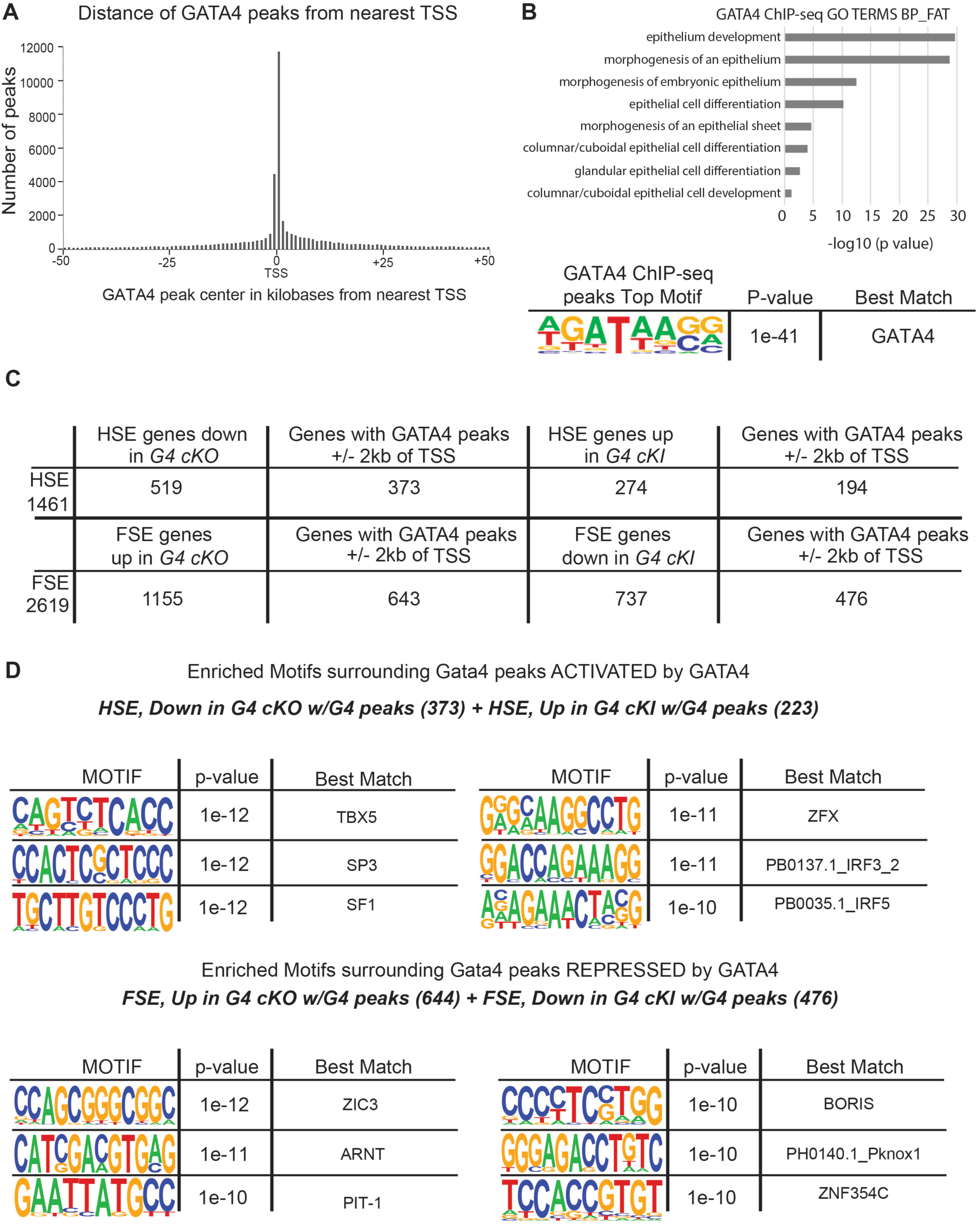
Putative GATA4 direct targets identified by cross-referencing RNA-Seq and ChIP-Seq datasets. (A) GATA4 ChIP-seq was performed with HS epithelial cells from adult mice. Peaks were mapped to the nearest transcriptional start site (TSS). The spatial distribution of GATA4 peaks within +/- 50 kb of the nearest TSS is shown. (B) DAVID analysis identified enriched gene ontology terms associated with peaks shown in (A). The GATA4 consensus sequence was the top motif present in peaks shown in (A), determined by *de novo* HOMER motif analysis. (C) GATA4 binding peaks from (A) were filtered to identify those within proximal regulatory regions of genes (+/- 2 kb of the TSS). This filtered peak set was cross-referenced with genes differentially regulated in *G4 cKO* and *G4 cKI* E14.5 epithelial cells and mapping to the HSE or FSE sets (Fig. 6B,C) to identify genes with HSE and FSE expression likely directly regulated by GATA4. (D) The *de novo* HOMER motif analysis tool was used to scan +/- 100 bp of the GATA4 binding peak center among genes identified in (C), predicted to be activated (HSE expression, down in HS mutants, up in FS mutants) or repressed by GATA4 (FSE expression, up in HS mutants, down in FS mutants). The top six enriched motifs are shown. Gene lists used can be found in supplemental table 5.

Of the genes with HSE expression down-regulated in E14.5 *G4 cKO* HS, 72% had GATA4 binding peaks within their proximal regulatory regions (Fig. 7C). Similarly, of the genes with HSE expression up-regulated in E14.5 *G4 cKI* FS, 71% had GATA4 binding peaks within their proximal regulatory regions (Fig. 7C). These represent genes hypothesized to be direct targets positively regulated by GATA4 such that expression is lost when GATA4 is deleted and induced when GATA4 is ectopically expressed. Looking at those genes we hypothesized to be direct targets negatively regulated by GATA4, such that expression is induced when GATA4 is deleted and lost when GATA4 is ectopically expressed, we found that expression of 56% of the FSE genes up-regulated in E14.5 *G4 cKO* HS and 65% of the FSE genes down-regulated in E14.5 *G4 cKI* FS contained GATA4 binding peaks within their proximal regulatory regions (Fig. 7C). These analyses allowed us to identify gene sets that we predict are directly induced or repressed by GATA4 in the developing stomach.

An outstanding question in GATA biology is the mechanism by which GATA factors function to activate or repress genes within a given tissue. To gain insight into this question in the context of gastric development, we examined the sequence neighborhoods surrounding GATA4 binding peaks within the proximal regulatory regions of genes positively or negatively regulated in *G4 cKO* HS and *G4 cKI* FS using HOMER *de novo* motif analysis. We identified different sets of consensus transcription factor binding motifs statistically enriched around GATA4 binding sites (+/- 100 bp of the peak) in the HSE and FSE gene sets differentially expressed in GATA4 mutants. Several *de novo* motifs predicted in the GATA4 binding neighborhoods of activated and repressed gene sets identified transcription factors previously shown to cooperate with GATA4 to direct gene regulatory networks in other organ systems including TBX5, SF1, and SP3 (Maitra et al. 2009; Nadeau et al. 2010; Ang et al. 2016; Watanabe et al. 2000; Tremblay and Viger 2001; Viger et al. 2008; Schteingart et al. 2019; Fokko van Loo et al. 2007). Others, although not yet definitively linked with GATA4, have transcriptional activities aligning with the GATA4 neighborhoods in which they were identified, i.e., positive regulators in activated genes or negative regulators in repressed genes (Rhie et al. 2018; Ni et al. 2020; Jheon et al. 2001; Kasaai et al. 2013; Watson, Tekki-Kessaris, and Boulter 2000; Longobardi et al. 2014; Maroni et al. 2017; Ciccarelli et al. 2016; Salgado-Albarrán et al. 2019; Scully et al. 2000; Sporici et al. 2005).

### GATA4 may regulate a transcription factor network to repress stratified squamous epithelial morphogenesis

To better understand the mechanism through which GATA4 promotes columnar epithelial cell development over stratified squamous epithelial cell development, we examined the status of known tissue-enriched transcription factors in GATA4 mutants. Previously, Sulahian et al. (2015) determined the expression of 1880 known and putative DNA-binding proteins in esophageal, gastric, and intestinal epithelial cells from 1-month old CD1 mice to identify those with tissue-enriched expression. This analysis yielded 21 transcription factors with enriched expression in esophagus versus stomach and intestine, and nine transcription factors with enriched expression in the stomach versus the intestine and esophagus. Three of the nine with gastric enriched expression and 15 of the 21 with esophageal enriched expression had similar enriched expression patterns in E14.5 HS or FS, respectively. All three gastric factors and 11 of the 15 esophageal factors contained experimentally validated GATA4 binding peaks within +/- 2kb of their TSS. Of these 3 gastric factors—ESRRG, PPARG, and NR0B2—only ESRRG was coordinately regulated in GATA4 mutants with its expression decreasing in *G4 cKO* HS epithelium and increasing in *G4 cKI* FS epithelium (Fig. 8A). With respect to the esophageal factors, 8 of the 11 with GATA4 binding peaks were coordinately regulated in GATA4 mutants with their expression decreasing in *G4 cKI* FS epithelium and increasing in *G4 cKO* HS epithelium (Fig. 8A). These results suggest a mechanism through which GATA4 promotes columnar fate in the HS epithelium by repressing expression of a network of stratified squamous enriched transcription factors.

**Figure 8.**
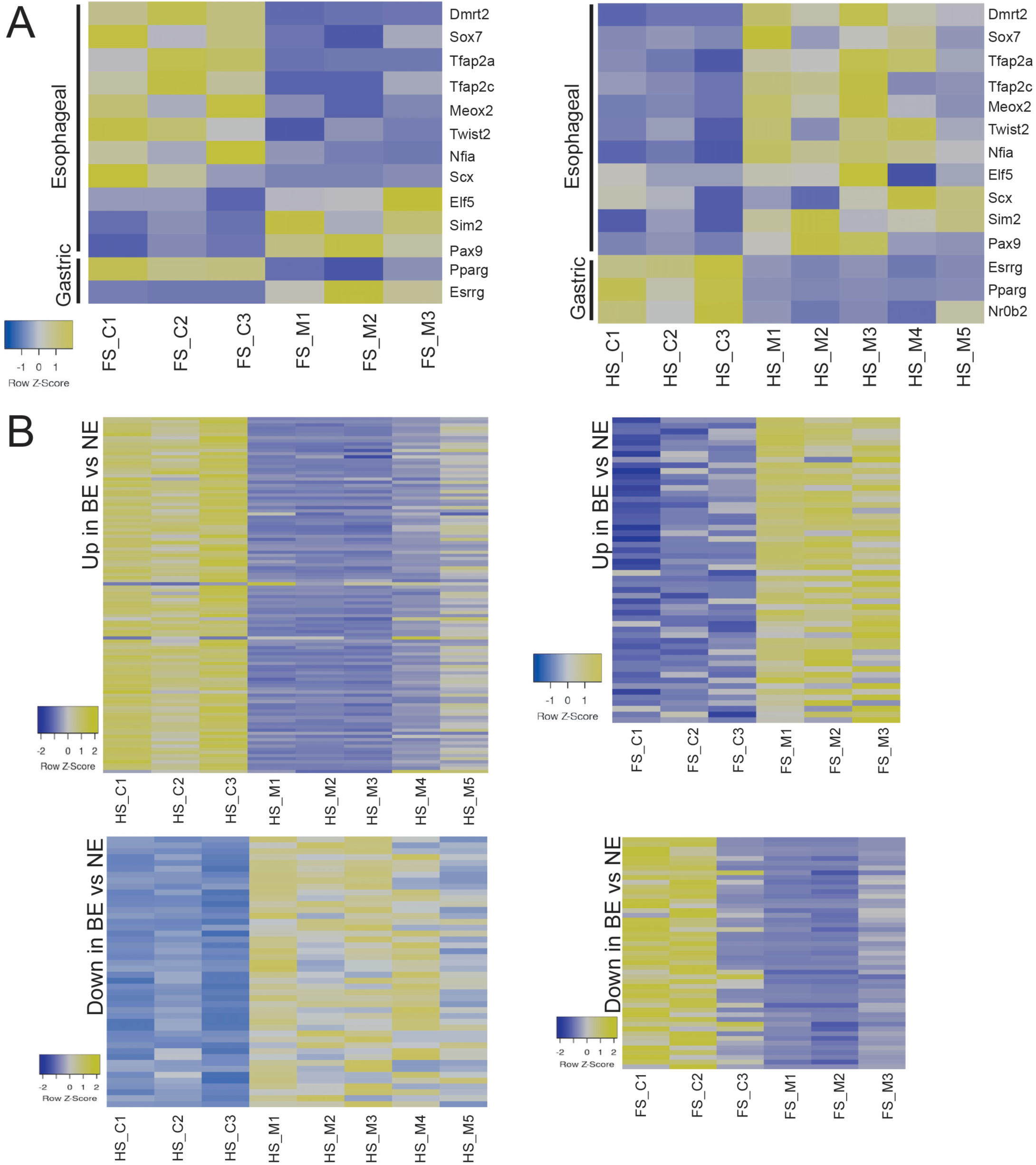
Transcriptional programs associated with esophageal/FS identity and human disease are altered in GATA4 mutant HS and FS epithelial cells. (A) Expression of a network of esophageal-specific transcription factors containing functional GATA4 binding sites were inversely regulated in *G4 cKO* and *G4 cKI* E14.5 HS and FS compared with controls. Heatmaps shown compare FPMK values for these factors between control and *G4 cKO* HS or control and *G4 cKI* FS. Esophageal transcription factors were induced in *G4 cKO* HS and suppressed in *G4 cKI* FS. Of a set of three gastric-specific transcription factors with functional GATA4 binding sites, expression of all was suppressed in *G4 cKO* HS, but only *Esrrg* was induced in *G4 cKI* FS. (B) Genes with differential expression in BE epithelium versus normal esophageal epithelium were cross-referenced with those genes differentially expressed in E14.5 *G4 cKO* HS or *G4 cKI* FS and containing functional GATA4 binding sites. Heatmaps compare FPMK values for BE-associated genes between control and *G4 cKO* HS or control and *G4 cKI* FS. Gene expression trends in GATA4 mutants correlated with trends in BE tissues. Genes induced in BE, i.e., columnar epithelial cell associated transcripts, were up-regulated in *G4 cKI* FS and down-regulated in *G4 cKO* HS. Genes suppressed in BE, i.e., stratified squamous epithelial cell associated transcripts, were up-regulated in *G4 cKO* HS and down-regulated in *G4 cKI* FS. Gene lists used for heatmap can be found in supplemental table 6.

### Genes regulated by GATA4 during gastric epithelial morphogenesis overlap with those dysregulated in Barrett’s esophagus

The columnar-like rather than stratified squamous epithelium observed in GATA4-expressing FS was reminiscent of BE, a premalignant stratified to columnar esophageal metaplasia. We and others have shown that GATA4 is abnormally expressed in BE (Miller et al. 2003; Haveri et al. 2008; Stavniichuk et al. 2020). To determine the extent to which altered gene expression patterns observed in GATA4-expressing FS epithelium and GATA4-deficient HS epithelium correlated with patterns observed in BE, we compiled a dataset of genes with known up- or down-regulated expression in BE versus normal esophageal tissue (Hyland et al. 2014; Sulahian et al. 2015; Zhang et al. 2019; Quante et al. 2012) and identified, among these BE-associated genes, GATA4 ChIP-Seq peaks in their proximal regulator regions. We examined gene expression patterns of these putative GATA4-regulated BE-associated genes in control HS, control FS, GATA4-expressing FS, and GATA4-deficient HS (Fig. 8B). As expected, the majority of BE-associated genes up-regulated in disease correlated with columnar epithelial cell profiles, being highly expressed in E14.5 HS controls and absent in FS controls. Expression of these genes was generally down-regulated in GATA4-deficient HS epithelium and induced in GATA4-expressing FS (Fig. 8B), paralleling the loss of columnar cell fate in mutant HS and the gain of columnar cell fate in mutant FS. The majority of BE-associated genes down-regulated in disease correlate with squamous epithelial cell profiles, being highly expressed in E14.5 FS controls and absent in HS controls. Expression of these genes was generally induced in GATA4-deficient HS epithelium and repressed in GATA4-expressing FS (Fig. 8B), paralleling the gain of stratified squamous cell fate in mutant HS and the loss of this profile in mutant FS (Fig. 8B). Among these putative GATA4-regulated BE-associated genes altered in GATA4 mutants, it was notable that the gastric and intestinal stem cell marker LGR5 was strongly induced in GATA4-expressing FS epithelial cells (+50-fold *G4 cKI*), suggesting a pivotal change in stem cell fate in mutant columnar-like FS. Taken together, these data correlate findings of a developmental model of GATA4 function in epithelial cell biology with a human disease in which GATA4 protein is mis-regulated.

## Discussion

During embryonic development, gene regulatory networks consisting of regionally restricted and lineage specific transcription factors regulate specification, morphogenesis, and differentiation of the GI tract epithelium. Our goal was to understand how GATA4, a key developmentally expressed transcription factor, participates in development and differential morphogenesis of the mouse stomach epithelium. Our studies using *Gata4* conditional knockout and knock-in mouse lines support the conclusion that GATA4 is necessary and sufficient for columnar epithelial morphogenesis in the stomach. Gastric epithelial morphogenesis was normal in *Gata6* conditional knockout embryos indicating that GATA4 uniquely controls this process. GATA4 must be expressed in the developing HS domain to permit columnar epithelial cell fate and repress stratified squamous epithelial cell fate. Conversely, GATA4 must be silenced in the developing FS domain to permit stratified squamous epithelial cell fate and block columnar epithelial cell fate. In this way, the establishment of domains with differential GATA4 expression in the developing mouse stomach determines gastric epithelial morphogenesis.

Our study agrees with and extends upon the previously published findings of Jacobsen et al. (2002) and Rodriguez-Seguel et al. (2020). Both studies provide a primarily histologic characterization of GATA4- dependent phenotypes in the developing stomach and show that glandular columnar epithelial architecture and mature HS cell types are abnormal in GATA4 mutant stomachs. Moving these studies forward, our study identified gastric defects earlier during development (E14.5 vs. E17.5-18.5), used both conditional knockout and knock-in models to demonstrate necessity and sufficiency of GATA4 in columnar HS development, and added in-depth global transcriptomic and DNA binding analyses to augment the understanding of GATA4’s role in gastric biology.

Simply identifying genes with up-regulated or down-regulated expression in GATA4 mutants, however, does not in itself indicate direct regulation of a gene’s transcription by GATA4. Identification of functional GATA4 binding sites within regulatory regions of differentially expressed genes can establish likely direct regulatory relationships between GATA4 and its putative targets. Overlapping expression data with GATA4 DNA binding data enabled us to establish connections between experimentally validated GATA4 binding sites and differentially expressed genes in our models to identify candidate genes most likely to be directly regulated by GATA4 in the developing murine stomach. One caveat to this analysis is that we used data from adult mouse HS epithelial cells rather than from E14.5 embryos. Although ideal to use embryonic cells, we encountered technical challenges. It is possible that our analyses have identified targets bound by GATA4 in the adult HS that aren’t similarly bound by GATA4 in the developing embryonic tissue (false positives) and that GATA4 binding peaks are missed because some targets are only bound by GATA4 during development (false negatives). Nevertheless, data from adult tissue can serve as a surrogate to demonstrate that GATA4 has the ability to bind to specific chromatin domains in simple columnar epithelium of the HS. Supporting the efficacy of using ChIP-Seq data from adult tissue to probe GATA4 function in embryonic tissue was the finding that GO terms for genes with GATA4 binding sites aligned with developmentally relevant biological functions including glandular columnar epithelial cell development, morphogenesis, and differentiation. Association of GATA4 binding sites with these themes further supports the idea that GATA4 is a critical regulator of columnar epithelial development in the glandular HS. The strong correlation between HSE and FSE genes differentially expressed in mutants and genes with experimentally validated GATA4 binding sites places GATA4 directly upstream of large networks of genes essential in mediating columnar HS development.

GATA factors generally, and GATA4 specifically, have been implicated as master regulators in other developmental systems. GATA4 activates expression of key proteins in a transcription factor network that guides cardiac development (Stefanovic and Christoffels 2015). In terms of repressive functions, GATA2/3 regulate a transcription factor network that promotes trophoblast differentiation by repressing pluripotency transcription factors in stem cells (Krendl et al. 2017). GATA1 promotes erythroid differentiation by repressing opposing upstream transcriptional activators of myelo-lymphoid differentiation while also suppressing expression of opposing factors’ downstream targets (Wontakal et al. 2012). These dual mechanisms of GATA1 function have a synergistic effect on lineage specification resulting in erythroid rather than myelo-lymphoid differentiation. The facts that functional GATA4 binding sites were identified in the promoters of more than half of a defined essential esophageal/forestomach enriched transcription factor set and that the expression of these transcription factors correlated with GATA4 presence or absence suggest that GATA4 directly represses expression of a transcription factor network in the developing HS to suppress stratified squamous epithelial development. Furthermore, downstream targets of these esophageal/forestomach transcription factors also contained GATA4 binding sites and changed expression state depending on the presence or absence of GATA4. Together, these data support the idea that GATA4 works through synergistic mechanisms to repress stratified squamous epithelial development in the HS.

Using information gained from cross-referencing of expression, chromatin binding, and motif analyses, we identified GATA4 co-factors that may contribute to GATA4’s ability to activate or repress transcription. Among these were several transcription factors that act with GATA factors to direct gene expression programs in other developmental systems including the heart, liver, and neural systems (Maitra et al. 2009; Nadeau et al. 2010; Ang et al. 2016; Watanabe et al. 2000; Tremblay and Viger 2001; Viger et al. 2008; Schteingart et al. 2019; Fokko van Loo et al. 2007). For example, TBX5 and GATA4 co-bind genes in cardiac cells to activate gene expression (Maitra et al. 2009; Nadeau et al. 2010; Ang et al. 2016). SF1 and GATA4 function cooperatively to activate gene expression pathways in Sertoli cells (Watanabe et al. 2000; Tremblay and Viger 2001; Viger et al. 2008; Schteingart et al. 2019). Although SP3, like GATA4, can function to activate or repress gene expression (Majello et al. 1997; Lania et al.1997), GATA4 and SP3 can interact to activate *Carp1* expression in the heart (Fokko van Loo et al. 2007). Because SP1 and SP3 bind the same consensus sequence (Lania et al. 1997), it is conceivable that SP1, typically a transactivating factor, serves as a GATA4 co-factor in the developing stomach. SP1 and GATA4 positively regulate cardiac and intestinal gene expression (Hu et al. 2011; von Salisch et al. 2011; Belaguli et al. 2007). Although not yet linked with GATA4, ZNF354C (ZFP354C) contains a KRAB transcriptional repression domain (Jheon et al. 2001; Kasaai et al. 2013; Watson et al. 2000). PKNOX1, which preferentially binds to promoter sequences, represses adipogenic differentiation and is implicated in negative regulation of the *GLUT4* gene (Longobardi et al. 2014; Maroni et al. 2017; Ciccarelli et al. 2016). BORIS negatively regulates an androgen receptor regulatory network in ovarian cancer (Salgado-Albarrán et al. 2019). Finally, PIT-1 functions with nuclear receptor co-repressor N-CoR to repress gene expression (Scully et al. 2000; Sporici et al. 2005). It is also possible that GATA4 binding to some promoters disrupts binding of other essential activating transcription factors to interfere with gene expression. We proposed this mechanism as a way GATA4 represses the *Fgf15* gene in the mouse jejunum (Thompson et al. 2017).

The morphological and molecular shifts from stratified to columnar and columnar to stratified epithelium observed in GATA4 mutant embryonic stomachs are reminiscent of tissue metaplasia (Giroux and Rustgi 2017). Stratified to columnar metaplasia is particularly relevant to the GI tract because Barrett’s metaplasia in the esophagus is a premalignant risk factor for EAC. Our previous work and the work of others have shown GATA4 to be aberrantly expressed in BE and EAC (Miller et al. 2003; Haveri et al. 2008; Stavniichuk et al. 2020). Our recent in vitro studies with human esophageal cell lines show that GATA4 expression responds to acid and bile and that, when over-expressed, GATA4 represses stratified marker gene expression including *Trp63*, *Krt5*, and *Krt15* (Stavniichuk et al. 2020). The correlation between dysregulated gene expression patterns in BE and GATA4 mutants as well as our finding that many of these genes have functional GATA4 binding sites support the idea that GATA4 does not merely serve as a marker of disease but that GATA4 plays a functional role in disease perhaps by supporting maintenance of columnar epithelium in BE. We propose that GATA4 regulates multiple BE associated pathways by activating or repressing the expression of essential columnar and stratified squamous cell associated transcription factors and their targets. For example, FOXA2, a columnar cell associated transcription factor up-regulated in BE, and TFAP2C, a stratified squamous cell associated transcription factor down-regulated in BE, both contain functional GATA4 binding sites and were mis-regulated in GATA4-expressing FS, with FOXA2 being induced and TFAP2C being repressed (Wang et al. 2014; Lv et al. 2019). These particular transcription factors were recently shown to be consistently altered across multiple BE expression studies with FOXA2 being activated and TFAP2C being suppressed in metaplastic cells (Lv et al. 2019). Although we propose that developmental studies like ours inform our understanding of human disease, we cannot with certainty conclude that GATA4 contributes to BE pathogenesis. Further studies using mice with GATA4 induction in the adult mouse esophagus or with alternative models such as human cell or organoid cultures from normal and BE tissues in which GATA4 is modulated will be crucial to understanding the role of GATA4 in BE.

## Acknowledgments

We thank Oscar Rosas Mejia and Rebekah Mokry for technical assistance. This work was supported by the National Institutes of Health (NIH), National Institute of Diabetes and Digestive and Kidney Diseases (DK111822, MAB) and the American Cancer Society (PF-19-023-01-DDC, AD).

## Author Contribution Statement

AD, BMK, ODF, RS, and CAT performed experiments, which were planned and supervised by MAB and SR. Data were analyzed by AD, BMK, KP, SR, and MAB. AD and MAB wrote the manuscript. Manuscript was reviewed by all authors.

## Competing interests

The author(s) declare no competing interests

## Data availability

The data discussed in this publication have been deposited in NCBI’s Gene Expression Omnibus and are accessible through GEO Series accession number GSE156324 (https://www.ncbi.nlm.nih.gov/geo/query/acc.cgi?acc=GSE156324).

## Notes

https://www.ncbi.nlm.nih.gov/geo/query/acc.cgi?acc=GSE156324

### Competing Interest Statement

The authors have declared no competing interest.

