## Supplemental Table 1 for "GATA4 regulates epithelial morphogenesis in the developing mouse stomach to promote establishment of a glandular columnar epithelium"

Supplementary Table 1: Genotyping primers

| Gene | Forward | Reverse | Size (bp) |
| --- | --- | --- | --- |
| *ShhCre* | gggacagctcacaagtcctc | ggtgcgctcctggacgta | 350 |
| *Gata4 cKO* alleles | | | |
| *Gata4^loxP^* | cccagtaaagaagtcagcacaaggaaac | agactattgatcccggagtgaacatt | 355, wt  455, loxP |
| *Gata4^null^* | ggggcaggacagcaagggggaggatt | gtgagacctgcagaatgggagtggagaatg | 207 |
| *Gata4 cKI* alleles | | | |
| *Rosa26^G4^* | aaagtcgctctgagttgttat | gcgaagagtttgtcctcaacc | 311 |
| *Rosa26^WT^* | aaagtcgctctgagttgttat | ggagcgggagaaatggatatg | 603 |
| *Gata6* *cKO* alleles | | | |
| *Gata6^loxP^* | gtggttgtaaggcggtttgt | acgcgagctccagaaaaagt | 159, wt  259, loxP |
| *Gata6^null^* | gctccaccctactatgaccaattcc | cccggtttaaaaatctgcttgagtc | 400 |
| *Gata4^flbio^* and *BirA* alleles | | | |
| *Gata4^Flbio^* | cagtgctgtctgctctgaagctgt | ccaaggtgggcttctctgtaagaa | 378, G4^wt^  550, G4^Flbio^ |
| *BirAJax14/15* | ttcagacactgcgtgact | ggctccaatgactatttgc | 500 (BirA+) |
| *BirAJax16/17* | gtgtaactgtggacagaggag | gaacttgatgtgtagaccagg | 400  (BirA-) |
