## Supplemental Table 2 for "GATA4 regulates epithelial morphogenesis in the developing mouse stomach to promote establishment of a glandular columnar epithelium"

Supplementary Table 2: Antibodies

| **Antibody** | **Dilution** | **Manufacturer** | **Catalog Number** |
| --- | --- | --- | --- |
| ATP4A  (mouse polyclonal) | 1:100 | Gift from Dr. Eunyoung Choi | n/a |
| GATA4  (rabbit monoclonal) | 1:500 | Cell Signaling  Danvers, MA | D3A3M |
| GATA4 C-20  (goat polyclonal) | 1:250 | Santa Cruz Biotechnology, Santa Cruz, CA | sc-1237 |
| GATA6  (rabbit monoclonal) | 1:500 | Cell Signaling  Danvers, MA | D61E4 |
| GIF  (goat polyclonal) | 1:2000 | Gift from Dr. Jason Mills | n/a |
| KRT5  (rabbit polyclonal) | 1:4000 | Biolegend (formerly Covance)  Dedham, MA | 905501 |
| KRT13  (rabbit monoclonal) | 1:250 | Abcam  Cambridge, UK | AB92551 |
| KRT14 (LL002)  (mouse monoclonal) | 1:1600 | Thermo Fisher  Rockford, IL | MA5-11599 |
| MUC5AC 45M1  (mouse monoclonal) | 1:100 | Thermo Fisher  Rockford, IL | MA1-38223 |
| TRP63  (rabbit polyclonal) | 1:250 | Abcam  Cambridge, UK | AB53039 |
| TRP63  (rabbit monoclonal) | 1:500 | Cell Signaling  Danvers, MA | D2K8X |
| Biotinylated horse-anti mouse IgG (H+L) | 1:66 | Vector Laboratories,  Burlingame, CA | BA-2000 |
| Biotinylated goat anti-rabbit IgG (H+L) | 1:66 | Vector Laboratories,  Burlingame, CA | BA-1000 |
| Biotinylated rabbit anti-goat IgG (H+L) | 1:66 | Vector Laboratories,  Burlingame, CA | BA-5000 |
