## Supplemental Table 3 for "GATA4 regulates epithelial morphogenesis in the developing mouse stomach to promote establishment of a glandular columnar epithelium"

Supplementary Table 3: TaqMan assay identifiers

| ***Gene*** | **TaqMan ID** |
| --- | --- |
| *Atp4a* | Mm00444417_m1 |
| *Cldn10* | Mm01226326_g1 |
| *Gata4* | Mm00484689_m1 |
| *Gata6* | Mm00802632_m1 |
| *Gif* | Mm00433596_m1 |
| *Hnf4a* | Mm00433964_m1 |
| *Krt15* | Mm00492972_m1 |
| *Krt17* | Mm00495207_m1 |
| *Krt23* | Mm00840789_m1 |
| *Mist1* | Mm00627532_s1 |
| *Muc5ac* | Mm01276718_m1 |
| *Muc6* | Mm00725165_m1 |
| *PepC* | Mm00482488_m1 |
| *Tff1* | Mm00436945_m1 |
| *Tff2* | Mm00447491_m1 |
| *Trp63* | Mm00495793_m1 |
| *Gapdh* | 4351309 |
